# Implications of the COVID-19 lockdown on dengue transmission in Malaysia

**DOI:** 10.1101/2020.07.21.214056

**Authors:** Song-Quan Ong, Hamdan Ahmad, Ahmad Mohiddin Mohd Ngesom

## Abstract

The impact of movement restrictions during the COVID-19 lockdown on the existing endemic infectious disease dengue fever has generated considerable research interest. We compared the Malaysia weekly epidemiological records of dengue incidences during the period of lockdown to the trend of previous years (2015 to 2019) and a simulation at the corresponding period that expected no movement restrictions. We found that the dengue incidence declined significantly with a greater magnitude at phase 1 of lockdown, with a negative gradient of 3.2-fold steeper than the trend observed in previous years and 6.5-fold steeper than the simulation, indicating that the control of population movement did reduce dengue transmission. However, starting from phase 2 of lockdown, the dengue incidences demonstrated an elevation and earlier rebound by at least 4 weeks and grew with an exponential pattern compared to the simulation and previous years. Together with our data on *Aedes* mosquitoes from a district of Penang, Malaysia, we revealed that *Aedes albopictus* is the predominant species for both indoor and outdoor environments. The abundance of the mosquito was increasing steadily during the period of lockdown, and demonstrated strong correlation with the locally reported dengue incidences; therefore, we proposed the possible diffusive effect of vector that led to a higher acceleration of incidence rate. These findings would help authorities review the direction and efforts of the vector control strategy.

## Introduction

Lockdown has been commonly implemented all over the world to halt COVID-19 transmission [1]. This is to control human movement, as physical proximity is a key risk factor for the transmission of SAR-CoV-2 [2]. Although the physical proximity of humans shapes the spatial spread of a pathogen, it refers mostly to directly transmitted infectious diseases [3] such as COVID-19, but the effect on indirectly transmitted infectious diseases such as dengue fever (DF) remains unclear.

Dengue fever (DF) is the most prevalent mosquito-borne disease in the world [4], understanding the impacts of human (host) movement on the transmission can improve the prevention system. Falcón-Lezama et al. [5] used a mathematical model to evaluate the effect of people’s day-to-day movement on the dengue epidemic and concluded that the vector-host’s spatial connectivity posted epidemic risk. To simulate the actual situation, Stoddard et al. [6] used contact-site cluster investigations in a case-control design to review the risk of dengue infection by human movement and argued the importance of movement restriction in managing the spatiotemporal dynamics of dengue virus. However, the previous experimental configurations were far from the real situation, especially when the mobility of the population on a large scale is not feasible to demonstrate the direct effect on dengue transmission. With the imposition of the COVID-19 partial lockdown in Malaysia on 13^th^ to 20^th^ weeks of 2020 (18 March to 12 May 2020), about 90% of people were restricted to their homes, and 10% essential workers were allowed to carry out their daily activities for the whole country [7]; thus, the unprecedented large-scale movement restriction of host provides opportunities to evaluate its direct impact on the dengue transmission.

To understand the direct impact of the large-scale movement restriction on Malaysia dengue endemic, comparison between the actual incidences trend with a simulation trend that expect no interference could be one of the approach as demonstrated by other studies on infection diseases [1–10]. To construct a best-fitting simulation, we have to understand the dengue endemic pattern in Malaysia, in which the dengue endemic in Malaysia is seasonal, with variable transmission and prevalence patterns affected by the large diversity in rainfall and spatial variation [11]. The major transmission periods of DF occur from June to September, following the main rainy seasons; the minor transmission period is from September to March, following monsoon rains sessions that bring higher precipitation and lead to the greater potential breeding ground for the vector [12]. The end of the minor peak and the start of major peaks of DF transmission often coincides with the duration of the COVID-19 lockdown (Figure 1). We postulate the trend of dengue incidences could be altered due to the changes of host movements, and seasonal time series analysis such as SARIMA is a suitable approach to construct the simulation as the method usually used to predict or forecast the infection incidences that were seasonal [8–10].

**Fig 1.**
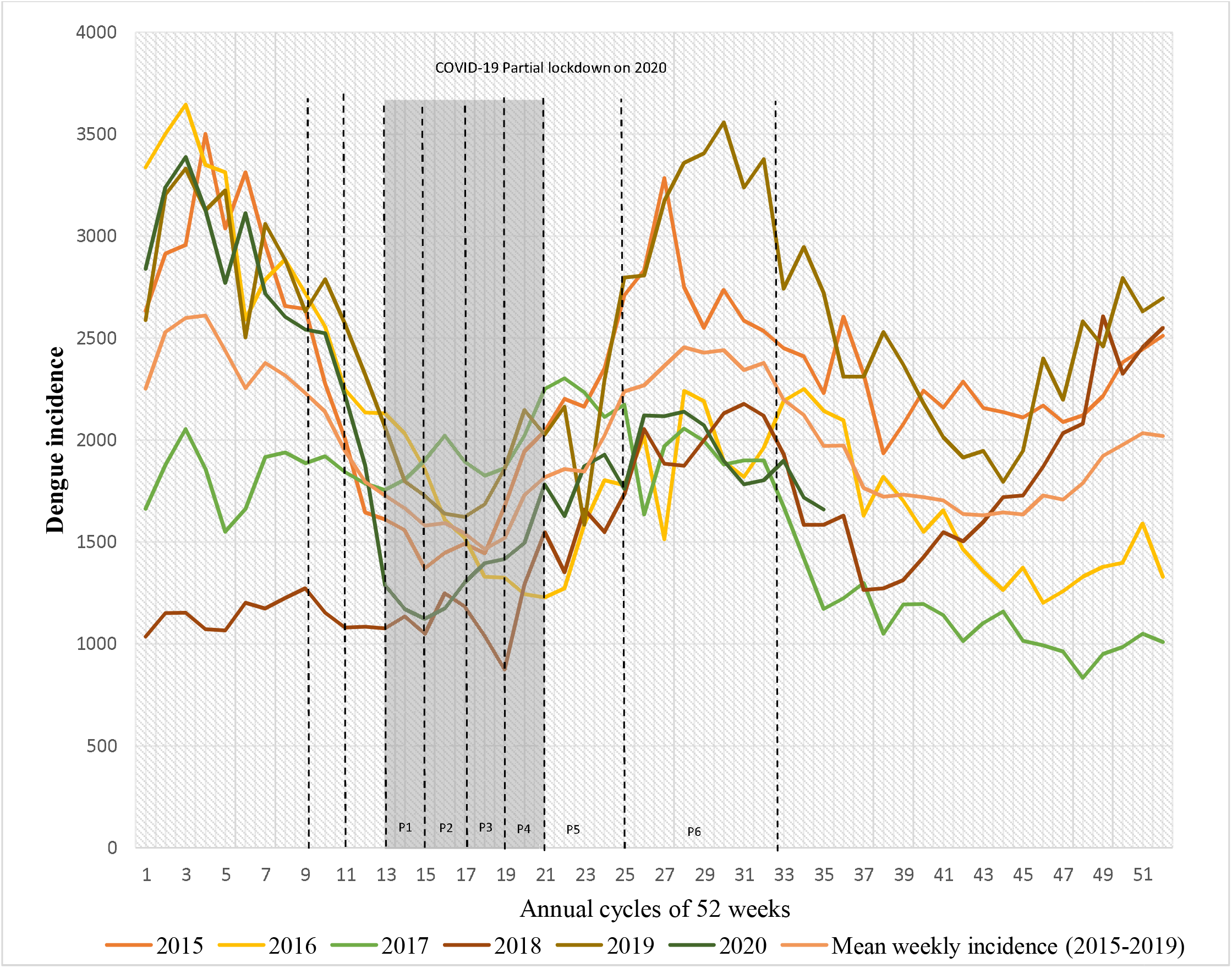
Dengue endemic growth from years 2015 to 2020. Eight phases: Pre2, Pre-lockdown 2; Pre1, Pre-lockdown 1; P1, Phase 1; P2, Phase 2; P3, Phase 3; P4, Phase 4; P5, Phase 5; P6, Phase 6

One of the key element in dengue transmission is the abundance and distribution of the vectors - *Aedes aegypti* and *Aedes albopictus*, in which the spatial distribution is also potentially affected by movement restrictions of humans. By using an agent-based transmission model, Reiner et al. [13] indicated that the socially structured movement of humans caused a significant influence on dengue transmission, and the infection dynamics were hidden by the diffusive effect of the vectors. The implementation of lockdown in Malaysia changed the host’s daily activities, most of the time people were contained in their own housing area, and also to both *Ae. aegypti* and *Ae. albopictus* that are highly anthropophilic, in which *Ae. aegypti* almost exclusively rely on human blood and *Ae. albopictus* is an aggressive and highly adaptive species that can easily colonize the habitat of other mosquitoes in urban areas [14]. In addition, the spatial distribution of the host also influenced the behavior of the vectors, and previous studies [14–18] identified a shift of the *Ae. albopictus* habitat to an indoor environment where the species usually inhabit the forest or are mostly vegetative and cause interspecies competition with other existing mosquito species, such as *Ae. aegypti*. Therefore, when the COVID-19 partial lockdown restricts humans in mostly indoor environments with minimum outdoor activities, we are interested in the distribution of the mosquitoes as well.

We study the effect of the physical proximity restriction on humans during the COVID-19 lockdown in Malaysia on two variables as shown in the paradigm in Fig. 2.

**Figure 2.**
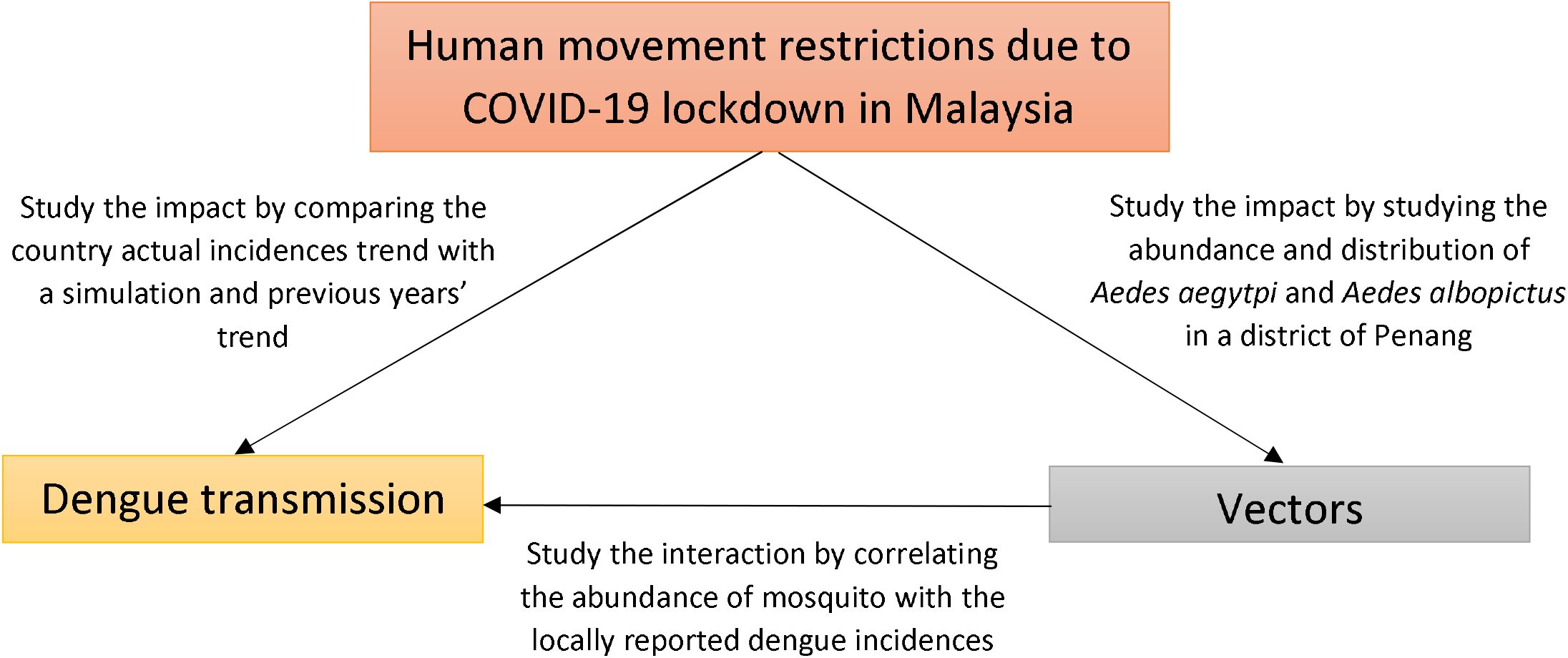
Paradigm of the study

### Country dengue incidences trend

This study aims to understand the impact of movement restrictions on the dengue transmission by statistically comparing the trend of actual reported dengue incidences with a SARIMA model simulation and previous years of incidences. We establish the SARIMA model simulation that expects no MR practice from the dataset of Malaysia weekly reported dengue incidences of previous years (2015 to 2019). We evaluate the level of heterogeneity of actual trend from simulation and previous years according to the phases of lockdown that implemented by Malaysia Government.

### The occurrence of vectors and correlation with locally reported incidences

To rationale the changes of country dengue incidences trend, we aim to obtain data on the abundance and distribution of *Aedes* mosquitoes during the period of lockdown by conducting sampling in the indoor and outdoor environments of areas from a district of Penang Malaysia.

## Methods

### Data collection

We started the analysis from the data of 2015, which recorded the second-highest dengue incidence after 2019 [19]. We retrieved the data from the official press statement and the Dengue Surveillance System developed by the Vector-Borne Disease Section, Ministry of Health (MOH), Malaysia, to monitor dengue transmission using remote sensing (RS) – iDengue [20], under the supervision of Remote Sensing Agency Malaysia. This surveillance system monitors dengue transmission across the country by updating dengue fevers reported in every hospital and medical institution on a daily basis, and all notified cases were followed up by the relevant health authorities for case verification before being recorded in the registry of the Dengue Surveillance System.

### Temporal analysis of Malaysia dengue incidences during partial lockdown

We observed the previous temporal pattern of dengue incidences and found that the period of the COVID-19 partial lockdown [13^th^ week (March 18) to 24^th^ week (June 9) of 2020] coincided between the end of minor (March-May) and the start of major (June-Sept) fluctuations of dengue transmission (Fig. 1). Therefore, to understand the heterogeneity of the trend of Malaysia weekly incidences due to the partial lockdown, we compared the actual trend with three reference trends, namely, a simulation, mean weekly incidence of year 2015- 2019, and the years that recorded the highest incidences trend (2019). The simulation was assumed to be without the interference of city lockdown and population movement control. To construct a simulation, we applied seasonal autoregressive integrated moving average (SARIMA) models, which are advantageous for modeling the seasonal and time-based dependent configuration of a time series and are commonly applied for epidemiological surveillance [8-10, 21]. We trained the models by using Malaysia weekly dengue incidences from 1^st^ week of 2015 to 52^nd^ week of 2019 from the dataset, and forecasted the incidences on 1^st^ to 35^th^ weeks of 2020, including the phases of partial lockdown in Malaysia (Table 1). We constructed and selected the best SARIMA model (p,d,q) x (P,D,Q) [p is the autoregressive lags, d is the degree of differencing, q is the moving-average lags, P is the seasonal autoregressive lags, D is the seasonal degree of differencing and Q is the seasonal moving-average lags] according to the methods of Box and Jenkins [22]. In addition, to obtain the best-fitting model, each of the constructed SARIMA model of forecast incidences before lockdown (1^st^ to 12^th^ weeks of 2020) was correlated with the actual weekly trend of dengue incidences by using Spearman rank correlation at the significance level of 0.05 (SPSS 17.0, IBM Corp.). The best-fitting model was selected based on the lowest values of the normalized Bayesian information criterion (NBIC) and the root mean square error (RMSE) [20], and the strongest correlation coefficient with the actual trend of 1^st^ to 12^th^ weeks of dengue incidences.

**Table 1.** Division of Eight Phases of Time Series for Data Analysis

| Phases | Terms | Time (Year 2020) | Corresponding weeks |
| --- | --- | --- | --- |
| Pre-2 | - | 19 February to 3 March | 8<sup>th</sup> to 9<sup>th</sup> |
| Pre-1 | - | 4 March to 17 March | 10<sup>th</sup> to 11<sup>th</sup> |
| P1* | MCO | 18 March to 31 March | 13<sup>th</sup> to 14<sup>th</sup> |
| P2 | MCO | 1 April to 14 April | 15<sup>th</sup> to 16<sup>th</sup> |
| P3 | MCO | 15 April to 28 April | 17<sup>th</sup> to 18<sup>th</sup> |
| P4 | MCO | 29 April to 12 May | 19<sup>th</sup> to 20<sup>th</sup> |
| P5 | CMCO | 13 May to 9 June | 21<sup>th</sup> to 24<sup>th</sup> |
| P6 | RMCO1 | 10 June to 31 August | 25<sup>th</sup> to 35<sup>th</sup> |
| P7 | RMCO2 | 1 September to 31 Dec | 36<sup>th</sup> to 52<sup>nd</sup> |

MCO - Movement Control Order
CMCO - Conditional Movement Control Order
RMCO - Recover Movement Control Order
\*for 2020, week 13 remarks the starting of COVID-19 partial lockdown

Furthermore, we divided the time series of 1^st^ to 35^th^ weeks of 2015-2020 into eight stages (two prelockdown periods, six phases during the partial lockdown) according to the announcement from the Malaysia government [23] (Table 1) and compared the actual weekly dengue incidences during the period of partial lockdown with the those of the simulation and previous years (2015-2019) using two-way ANOVA with two independent variables, namely, trends (actual/simulation/previous years of incidences), and phases. Because the trends follow an open-up parabolic pattern, we further distinguish the pattern by comparing the slopes for the phases (based on incidences in three weeks) to study the rates of decline and endemic incline.

### Mosquito occurrences

We assessed the abundance and distribution of *Aedes* mosquitoes during the movement restriction of human in particular the indoor and outdoor environments due to the possibility of a decrease in artificial breeding sites reported at the beginning of lockdown [24] and the host contained in their housing area. The sites were selected from three residential areas, namely, Taman Bukit Jambul (5°20’06.6”N 100°17’18.7”E), Taman Permai Indah (5°20’54.4”N 100°17’52.9”E) and Taman Jelutong (5°23’18.3”N 100°18’37.7”E) which are located within the Northeast Penang Island District (Fig. 3) and were the dengue hotspot area [25]. Due to the restriction of movement order from Malaysia government [26], the sampling was conducted at limited coverage of area, with five outdoor (n=5) and indoor (n=5) locations from each of the sampling area, respectively. For high anthropophilic mosquitoes, the human landing catch (HLC) method is the most effective sampling method [27], although it poses the risks of the human contracting the mosquito-borne pathogen, especially at the location where dengue and chikungunya are endemic. Three participants were voluntarily participated to the experiments and all their informed consents were confirmed and obtained prior to sampling. All experimental methodologies were approved by the ethical committee of UOW Malaysia KDU Penang University College and in accordance with the guidelines of the ethical committee of UOW Malaysia KDU Penang University College. The caught was performed in the early morning from 7:30 A.M. to 10 A.M. with the left arm and both legs exposed without any artificial chemical (e.g., lotion and body shampoo) interference; the mosquitoes were collected by a manual aspirator and transferred to a sealed container. Mosquitoes were killed by freezing, counted, and identified using taxonomy keys. The identification of mosquitoes was based on the distinguish features - the lyre-shaped markings on *Ae. aegypti* and the white stripe marking on the thorax of *Ae. albopictus*, we also focused on the clypeus-pedicel parts and mesepimeron of *Ae. aegypti* that consisted of distinctive white scales. The counts of *Ae. aegypti* and *Ae. albopictus* from indoor and outdoor locations for the eight stages of the time series during partial lockdown were obtained and correlated with the weekly reported dengue incidences of Penang, Malaysia by using Spearman rank correlation at the significance level of 0.05 (SPSS 17.0, IBM Corp.).

**Figure 3.**
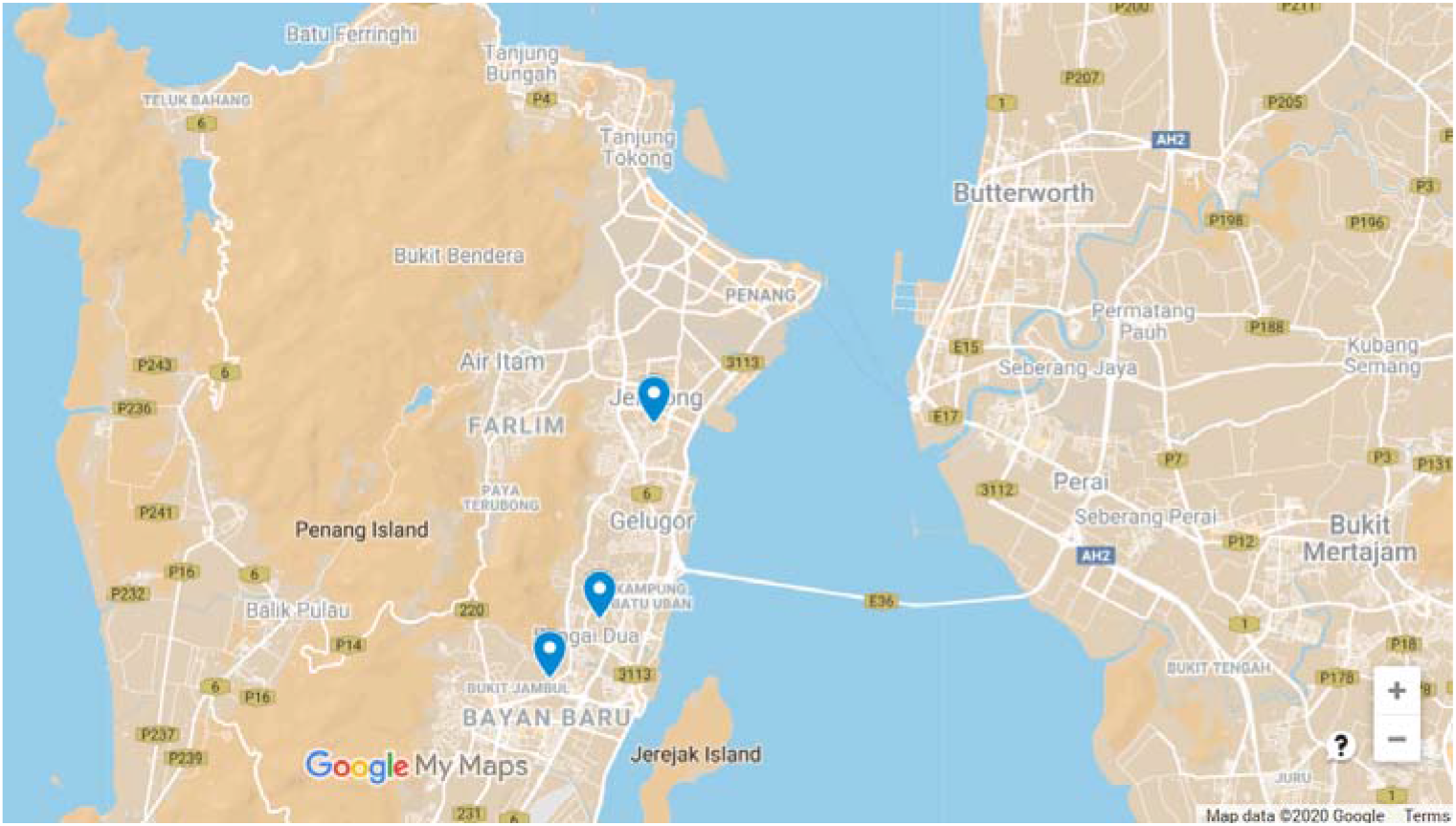
Sampling locations of *Aedes* mosquitoes

## Results

### Temporal analysis of dengue incidences during partial lockdown

Most of the studies on the growth of dengue incidences in Malaysia have focused on the total cases reported annually, in which the proposed model may not be sensitive and flexible enough to predict growth and propose necessary management. To our knowledge, this study is the first to report a simulation model with weekly dengue incidences in 2020, including the period of COVID-19 partial lockdown in Malaysia. To select the best-fitting simulation model, Table 2 shows the comparison of the NBIC, RMSE, and MAPE of nine SARIMA models, and the best-fitting forecast model for dengue incidences is SARIMA (1,1,0) (1,1,1) model due having the lowest NBIC and RMSE, strongest correlation (r=0.718, p<0.05) between the forecasted and actual incidences trend before partial lockdown (1^st^ to 12^th^ weeks of 2020), and the model also passed the Ljung–Box Q Test (z = 19.782, p = 0.180).

**Table 2.**
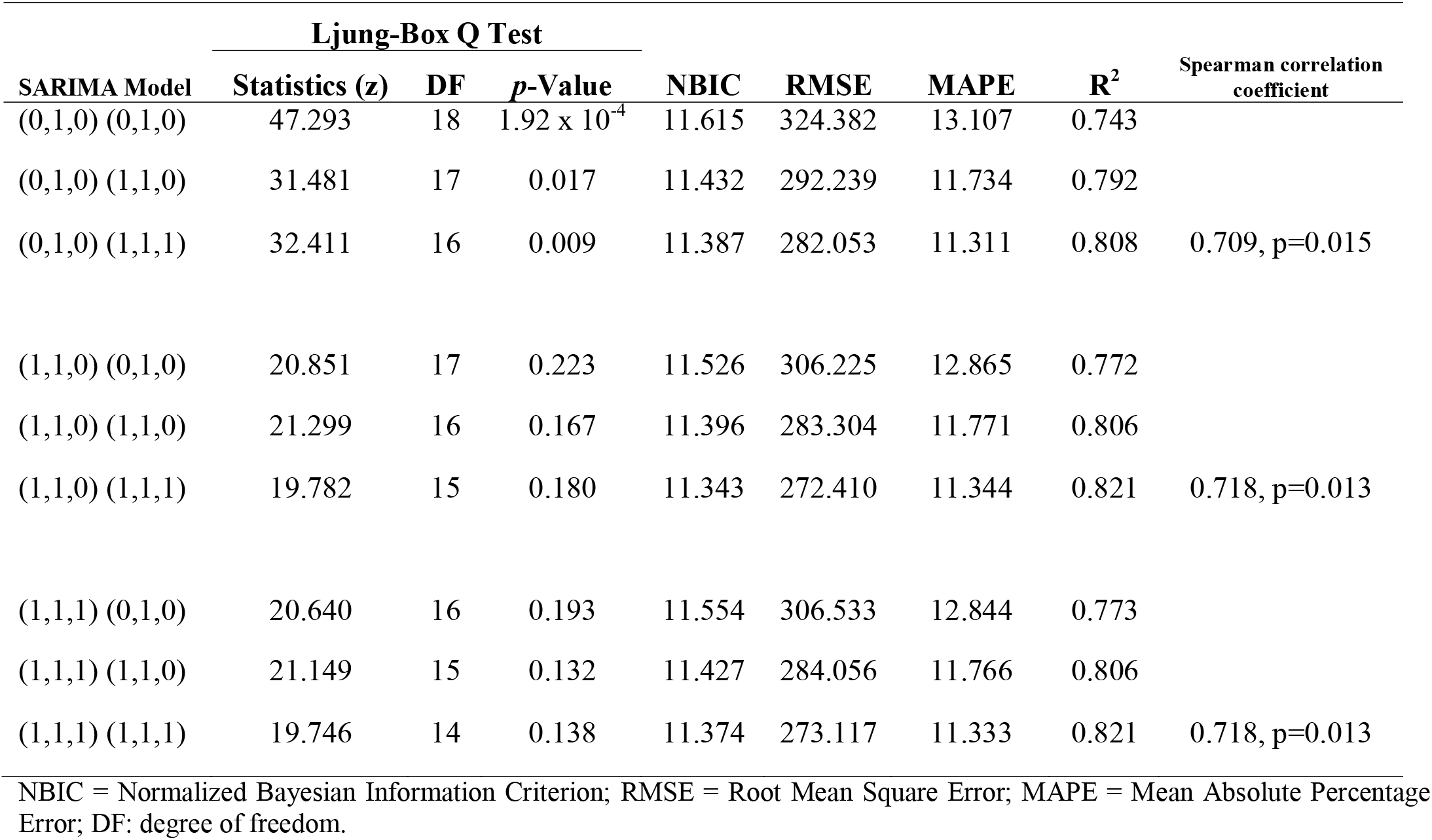
Comparison of candidate SARIMA models.

As seen in Fig. 4, when we generated a simulated model on the trend of 13^th^ to 35 weeks of 2020, which coincided with the COVID-19 partial lockdown period in Malaysia, the actual dengue incidence trend was significantly diverted and demonstrated a stronger negative steepness compared with the simulated trend. The actual trend is lower than the lower confidence level of forecasted trend (Fig.4), which implied that the actual dengue incidence trend significantly disobeyed the simulation, which presumed no lockdown, indicating that movement control greatly impacted dengue transmission in Malaysia. To further analyze the changes in dengue incidence trends due to the partial lockdown, we divided the time series of endemical weeks into eight stages as described in Table 1, and Fig. 5 describes the ANOVA comparison and Table 3 showed the slopes of the eight stages for previous years (mean weekly incidence of 2015-2019), the SARIMA model simulated trends, year 2019 and actual dengue incidences during the COVID-19 partial lockdown. Some researchers have proposed that the dengue incidences in 2020 were lower than those in 2019 [16], but when we averaged the weekly incidences of the previous five years (2015-2019), 2020 had significantly higher dengue incidences at prelockdown 1 and 2 even though the slopes between the dengue incidences of the previous years and those of the year 2020 during the period of prelockdown (1 and 2) are fairly the same. When Malaysia imposed phase 1 of the partial lockdown, the slope declined dramatically, which was 319% steeper than in previous years and 650% steeper than SARIMA simulated trend. These provide a strong implication that movement control during partial lockdown significantly reduced the reported dengue incidences. Although at phases 2 to 4, the incidences in 2020 were significantly lower than those in previous years, when we compared the stages for the slope to become positive (which indicates an upsurge in dengue incidences), this change in slope occurred in 2020 two stages (4 weeks) earlier than in previous years; specifically, a positive slope was obtained at phase 3 in 2020 compared to in previous years, in which a positive was obtained at phase 5. Furthermore, at phase 5, the steepness of the slope of the year 2020 spiked from phase 4 to phase 5 compared to previous years, with the steepness increasing from 22% to 227%, suggesting a significant increase in dengue transmission.

**Figure 4.**
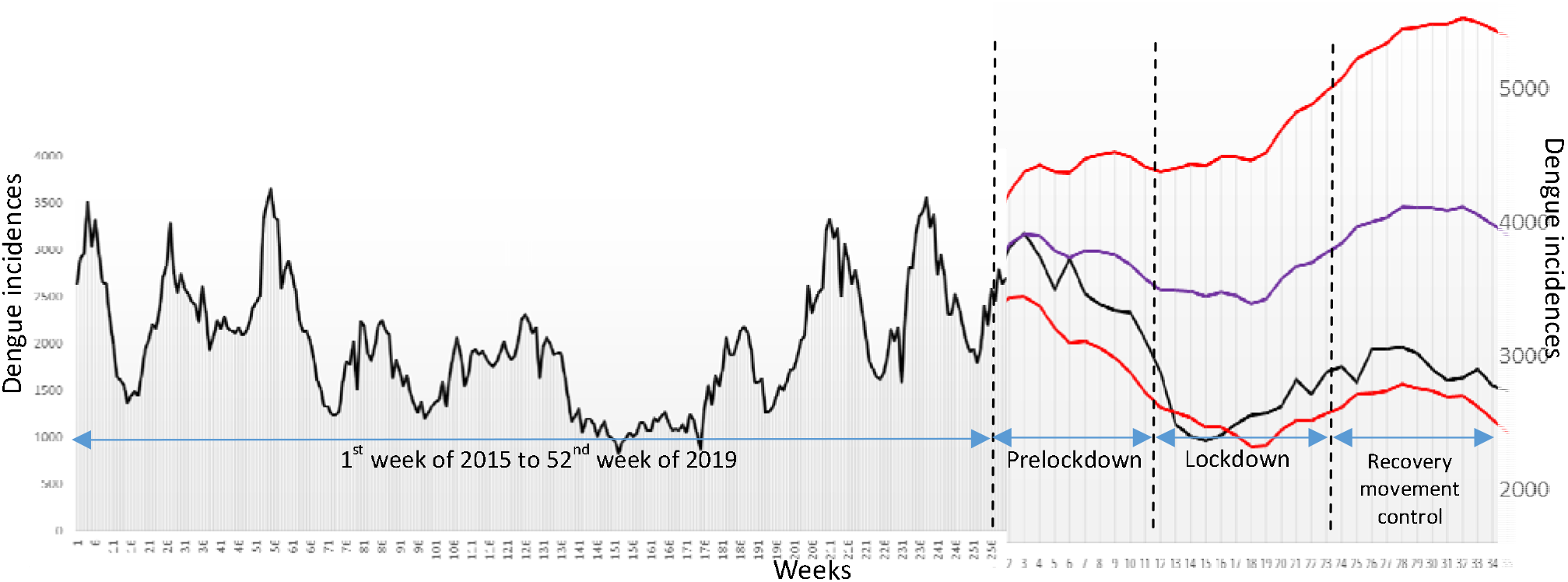
Comparison between actual weekly dengue incidence from 1^st^ week of 2015 to 35^th^ of 2020 with SARIMA model forecasted simulation before and the duration of partial lockdown. ULC, upper confidence level; LCL, lover confidence level.

**Figure 5.**
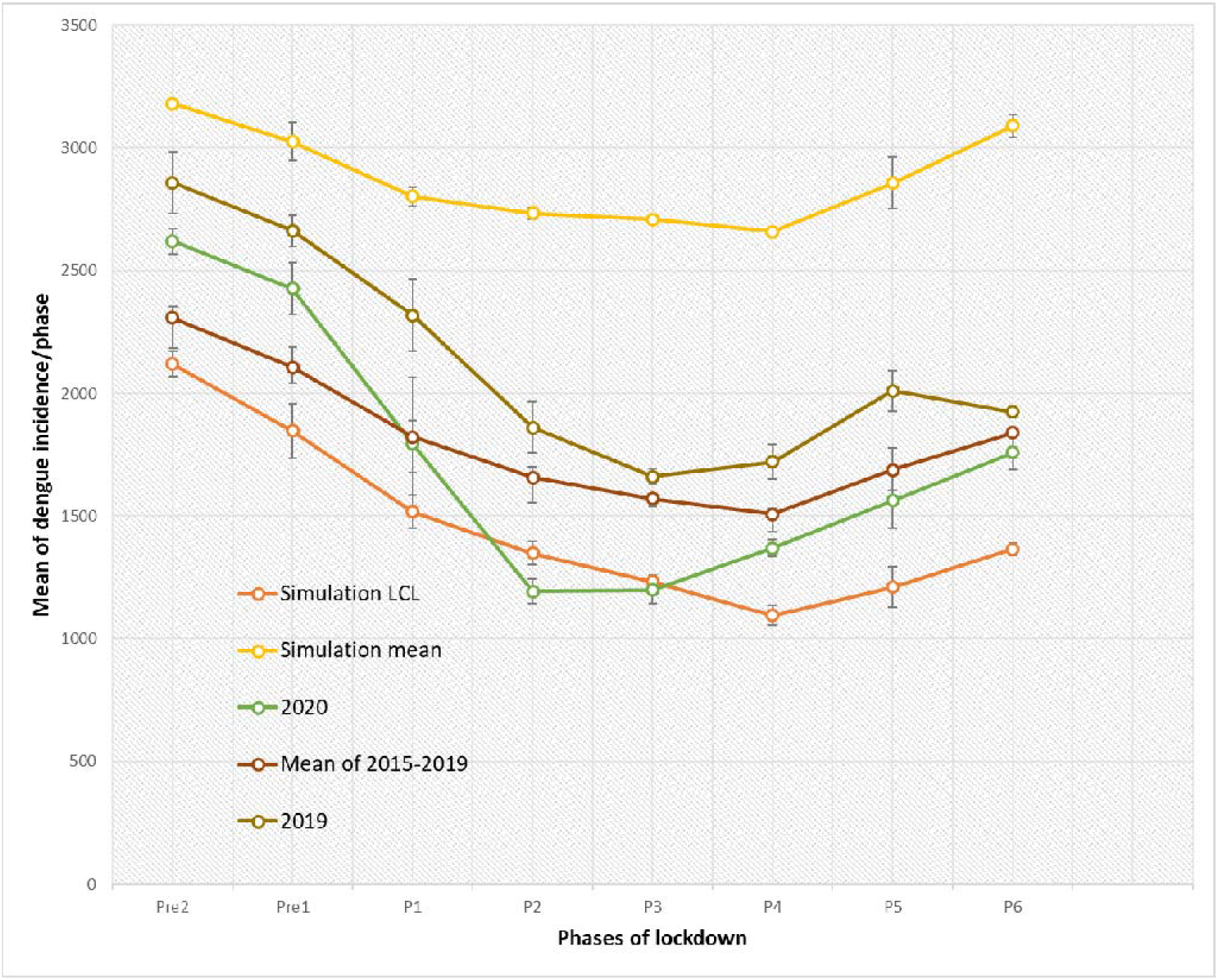
Analysis of variance (ANOVA) of dengue incidences of mean, simulation (SARIMA model) and year 2020 for eight phases of COVID-19 partial lockdown in Malaysia. Pre - Pre-lockdown; P - phase

**Table 3.** Slope for the phases

|  | Slope |  |  |  |  |  |  |  |
| --- | --- | --- | --- | --- | --- | --- | --- | --- |
|  | Pre2 | Pre1 | P1 | P2 | P3 | P4 | P5 | P6 |
| Simulation mean | -25.60 | -134.62 | -62.11 | -33.75 | 4.96 | -18.36 | 182.13 | 74.75 |
| 2020 | -88 | -161 | -466 | -83 | 90 | 57 | 183.5 | 44.5 |
| Mean of 2015-2019 | -74.1 | -140.6 | -111.2 | -73.2 | -19 | -11.9 | 149.5 | 13.6 |
| 2019 | -214 | -31 | -253 | -169 | -51.5 | 118 | 83.5 | -221 |

Pre - Pre-lockdown; P - phase

### Mosquito occurrences

To study one of the factors that contributes to the spread-out of dengue transmission during the COVID-19 partial lockdown, we assessed the abundance and distribution of vectors during the partial breakdown. Fig. 6 shows the total numbers of *Ae. albopictus* collected from the outdoor area of the sampling locations during the period of partial lockdown. *Ae. albopictus* is the predominant species in the outdoor area, with no *Ae. aegypti* was caught. The abundance of *Ae. albopictus* showed slight fluctuation patterns during the partial lockdown but still demonstrated a strong linear increment throughout the eight phases of partial lockdown, and showed strong correlation with Penang reported dengue incidences (r=0.952, p<0.001). In general, the total number of mosquitoes caught indoors was significantly lower than that outdoors, and both *Ae. aegypti* and *Ae. albopictus* were caught indoors, with the abundance of *Ae. aegypti* being relatively low and plateauing throughout phase 3 to 6. However, *Ae. albopictus* demonstrated higher abundance and exponential growth with the population during the same corresponding period (Table 4).

**Figure 6.**
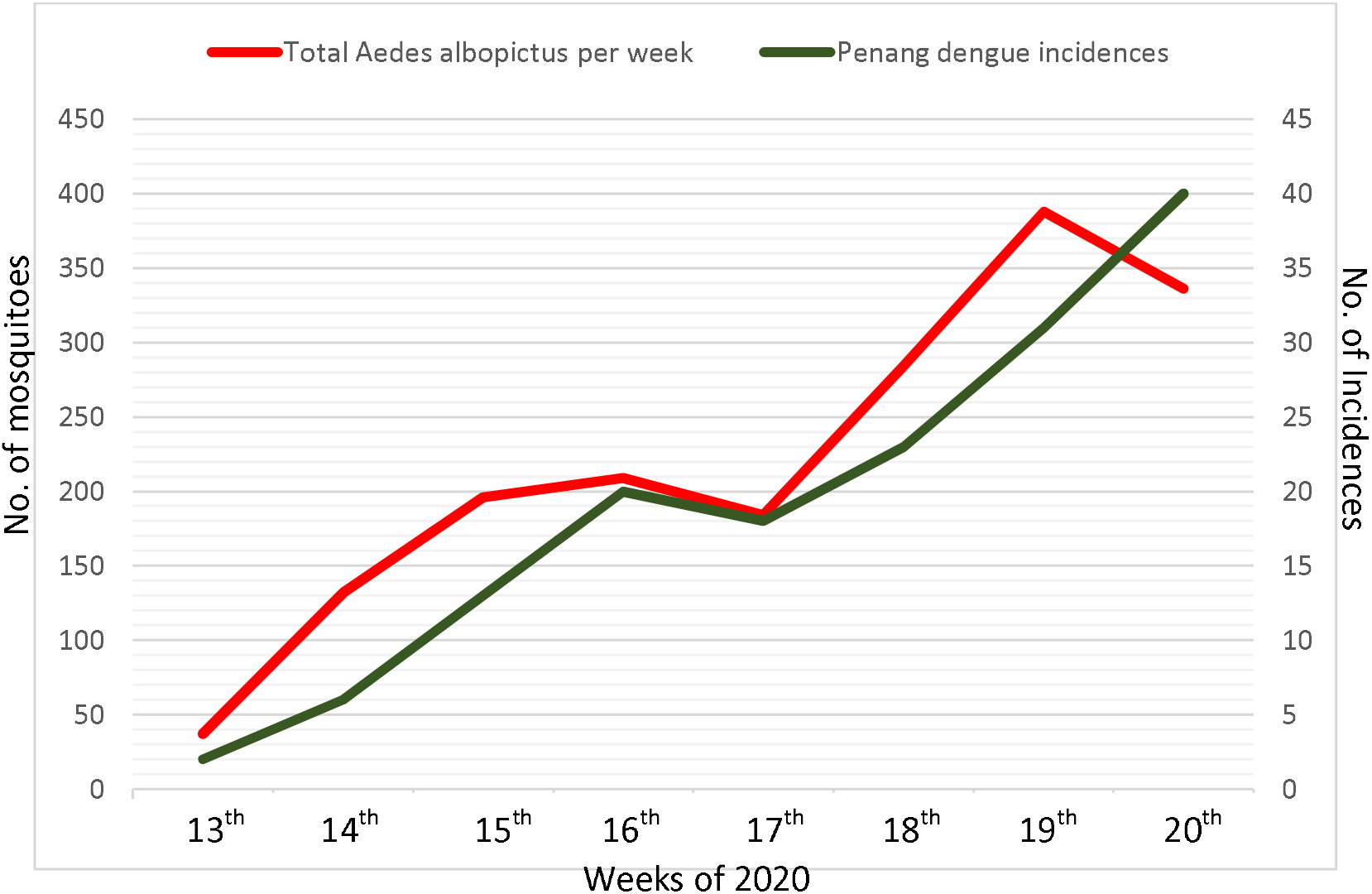
Total counts of Aedes albopictus for the weeks and phases of COVID-19 partial lockdown at the outdoor environment of sampling locations.

**Table 4.** Total *Aedes* mosquitoes caught from indoor environment of sampling locations

| Phases | *Aedes aegypti* | *Aedes albopictus* |
| --- | --- | --- |
| P3 | 0 | 4 |
| P4 | 3 | 2 |
| P5 | 4 | 9 |
| P6 | 2 | 12 |

P - Phase

## Discussion

The factors that contribute to dengue transmission are multifaceted, and the spatial variation in the contact rates of the host and vector are probably the most important factors for the dynamics of DENV [28]. With the movement restrictions (MRs) imposed in Malaysia due to the COVID-19 pandemic, we can investigate the effect of the large-scale MRs of the host on two interrelated variables: dengue transmission and *Aedes* mosquitoes occurrences. We analyze the dengue incidence trends by comparing their significant differences among the phases before and during the lockdown to those of the same corresponding periods for previous years and simulation. We first reported evidence that the MRs of the COVID-19 partial lockdown significantly influenced the weekly dengue incidence trend in Malaysia. Our findings provide direct evidence from analysis and extend the studies of Reiner et al. [13] and Falcón-Lezama et al. [5], which demonstrated that people’s movement affected dengue transmission by using a simulation model. The early decline in dengue incidences was also reported in India, with dengue cases dropping by 50% compared to previous years. The decline of incidences at the beginning of the lockdown could have occurred for several reasons: (1) fewer hosts available outdoors and therefore less vector-host contact, as *Ae. aegypti* and *Ae. albopictus* are exophilic [29]; (2) the alteration of the environment and relatively fewer artificial breeding sites for the vector due to less solid waste from humans [24]; and (3) the limited movement of infected patients due to the COVID-19 partial lockdown.

Unfortunately, our analysis showed that the dengue incidences rebounded earlier and spiked up at a higher rate than in previous years, indicating that the large-scale MRs of the population are not sustainable in controlling the spread of dengue. The finding is compatible with the situation in Singapore, which has had the most serious dengue outbreak in seven years [30], and an agent-based simulation model study by Jindal and Rao [31] that showed a significantly higher risk and severity of dengue transmission after the COVID-19 partial lockdown. The stay-at-home situation makes the host available most of the time in the indoor environment and optimizes the biting activities for endophagic *Ae. aegypti* to transmit the virus. In contrast to Harrington et al. [32], who argued that people, rather than mosquitoes, rapidly move the virus within and between rural communities and places due to the limitation of the flight range of female *Ae. aegypti*, our result of *Aedes* mosquitoes revealed that the element of vector dispersal plays a more crucial role in spreading the virus, when the abundance growth of the sampling locations correlated strongly with locally reported dengue incidences. We also suggest that *Ae. albopictus* could be the key substitution vector that contributes significantly to dengue virus circulation, and therefore, the vector control direction and strategies should be redesigned.

With no specific previous entomological data about female adult *Ae. albopictus* in the corresponding period with lockdown, we refer to Rozilawati et al. [14] and Rahim et al. [33], who studied the seasonal abundance of *Ae. albopictus* in Penang by sampling eggs, the Ovitrap index, the container index (CI), the house index (HI) and the Breteau Index (BI). Their results demonstrated that the indexes of *Ae. albopictus* for the corresponding period of phase 2 to 4 of lockdown should be lower, in contrast to our finding that the abundance of *Ae. albopictus* increased steadily from phases 1 to 5. There are several reasons for the increase in *Ae. Albopictus*. First, as proposed by the WHO [34], the upsurge of *Aedes* mosquitoes may be due to the southwestern monsoon (end of May to September), which brought a higher frequency of precipitation and higher humidity and temperature, and therefore, a higher breeding rate for the mosquitoes. Second, a minimum centralized vector control program can be conducted due to the stay-at-home policy. It is relevant to any method that is intended to reduce dengue incidences by reducing, but not eliminating, *Aedes* mosquito populations. Before that, researchers [35] have associated the index of the temporal vector with dengue occurrence, and the relationships between *Aedes* mosquito density and DENV transmission indexes for *Ae. aegypti* density are correlated with the prevalence of human dengue infections but are relatively weakly correlated with the incidences, indicating that other factors were involved in determining the incidence pattern. This is supported by the participant during the *Aedes* survey when a fogging activity was observed on May 28, 2020 (22^nd^ weeks of 2020, phase 5 – conditional movement control order), and total *Ae. albopictus* was significantly lower on May 29, 2020, but the mosquitoes caught afterward remained elevated in general.

Furthermore, our findings showed the presence of both *Ae. agypti* and *Ae. albopictus* from an indoor environment but no *Ae. aegypti* from the outdoors, indicating that *Ae. albopictus* is better adapted to a sudden change in the environment, such as the duration of lockdown when most of the hosts shift to the indoors. With the consistent growth rate of the indoor and outdoor populations, we postulate that *Ae. albopictus* invades the habitat of *Ae. aegypti* and showed a high possibility of colonizing the habitat. Our result is consistent with Nur Aida et al. [36] and Dieng et al. [18], who found that they could increase the invasiveness of *Ae. albopictus* by obtaining a high number of egg and mosquito counts from the indoor environment of Penang Island. Previous studies [14, 16, 37] from other countries have also reported the aggressive invasive behavior of *Ae. albopictus*, which shared the habitat with other native or existing mosquitoes, including *Ae. aegypti*, which are commonly predominant in indoor environments [38]. Due to the restriction of traveling during the period of lockdown, our results provided limited area coverage, but when considering the scale of the study as a districts-level assessment, the results propose a few important discoveries of vector distribution and occurrence during the MRs of lockdown.

## Acknowledgments

We extend our deepest gratitude to the Ministry of Health Malaysia for providing the data.

## References

1. Sault A. Why lockdowns can halt the spread of COVID-19. World Economic Forum 2020 March 21. Available from https://www.weforum.org/agenda/2020/03/why-lockdowns-work-epidemics-coronavirus-covid19/ Accessed 10 May 2020

2. World Health Organization (WHO). Modes of transmission of virus causing COVID-19: implications for IPC precaution recommendations. Available from: https://www.who.int/news-room/commentaries/detail/modes-of-transmission-of-virus-causing-covid-19-implications-for-ipc-precaution-recommendations Accessed 22 April 2020

3. Mossong J, Hens N, Jit M, Beutels P, Auranen K, Mikolajczyk R et al. Social contacts and mixing patterns relevant to the spread of infectious diseases. PLoS Medicine. 2008; 5 http://dx.doi.org/10.1371/journal.pmed.0050074.

4. World Health Organization (WHO). Dengue and severe dengue. Available from: https://www.who.int/news-room/fact-sheets/detail/dengue-and-severe-dengue Accessed 24 June 2020

5. Falcón-Lezama JA, Martínez-Vega RA, Kuri-Morales PA, Ramos-Castañeda J, & Adams B. Day-to-Day Population Movement and the Management of Dengue Epidemics. Bull. Math. Biol. 2016; 78(10), 2011–2033. https://doi.org/10.1007/s11538-016-0209-6

6. Stoddard ST, Forshey BM, Morrison AC, Paz-Soldan VA, Vazquez-Prokopec GM, Astete H et al. House-to-house human movement drives dengue virus transmission. Proceedings of the National Academy of Sciences of the United States of America, 110(3), 994–999. https://doi.org/10.1073/pnas.1213349110

7. Charles Hector. Operating businesses during MCO must be classified as offence. Available from: https://www.malaysiakini.com/letters/519902 Accessed 3 May 2020

8. Zhang X, Liu Y, Yang M, Zhang T, Young AA, Li X. Comparative study of four time series methods in forecasting typhoid fever incidence in China. PLoS One. 2013 May 1;8(5):e63116. doi: 10.1371/journal.pone.0063116. PMID: 23650546; PMCID: PMC3641111.

9. Martinez, Edson Zangiacomi, Silva Elisângela Aparecida Soares da, & Fabbro Amaury Lelis Dal. (2011). A SARIMA forecasting model to predict the number of cases of dengue in Campinas, State of São Paulo, Brazil. Revista da Sociedade Brasileira de Medicina Tropical, 44(4), 436–440. https://doi.org/10.1590/S0037-86822011000400007

10. Cong J, Ren M, Xie S, Wang P. Predicting Seasonal Influenza Based on SARIMA Model, in Mainland China from 2005 to 2018. Int J Environ Res Public Health. 2019 Nov 27;16(23):4760. doi: 10.3390/ijerph16234760. PMID: 31783697; PMCID: PMC6926639.

11. Hii YL, Zaki RA, Aghamohammadi N, Rocklöv J. Research on Climate and Dengue in Malaysia: A Systematic Review. Curr Environ Health Rep. 2016;3(1):81–90. doi:10.1007/s40572-016-0078-z

12. Sulaiman S., Pawanchee Z.A., Jeffery J., Ghauth I. & Busparani V. Studies on the distribution and abundance of *Aedes aegypti* (L.) and *Aedes albopictus* (Skuse) (Diptera: Culicidae)in an endemic area of dengue/ dengue hemorrhagic fever in Kuala Lumpur. Mosquito-Borne Diseases Bulletin 1991; 8: 35–39.

13. Reiner RC, Jr Steven T. Stoddard, and Thomas W. Scotta. Socially structured human movement shapes dengue transmission despite the diffusive effect of mosquito dispersal. Epidemics. 2014 March 6; 30–36. doi:10.1016/j.epidem.2013.12.003.

14. Li Y, Kamara F, Zhou G, Puthiyakunnon S, Li C, et al. Urbanization Increases *Aedes albopictus* Larval Habitats and Accelerates Mosquito Development and Survivorship. PLoS Negl Trop Dis. 2014; 8(11): e3301. doi:10.1371/journal.pntd.0003301

15. Rozilawati H, Zairi J, Adanan CR. Seasonal abundance of *Aedes albopictus* in selected urban and suburban areas in Penang, Malaysia. Trop Biomed. 2007;24(1):83–94.

16. Rodrigues Md, Marques GRAM, Serpa LN et al. Density of *Aedes aegypti* and *Aedes albopictus* and its association with number of residents and meteorological variables in the home environment of dengue endemic area, São Paulo, Brazil. Parasites Vectors 2015; 8, 115. https://doi.org/10.1186/s13071-015-0703-y

17. Estelle Martin, Matthew C.I. Medeiros, Ester Carbajal, Edwin Valdez, Jose G. Juarez, Selene Garcia-Luna et al. Surveillance of *Aedes aegypti* indoors and outdoors using Autocidal Gravid Ovitraps in South Texas during local transmission of Zika virus, 2016 to 2018. Acta Tropica. 2019;192: 129–137. https://doi.org/10.1016/j.actatropica.2019.02.006.

18. Dieng H, Saifur RGM, Hassan AA, Salmah MRC, Boots M, et al. Indoor-Breeding of *Aedes albopictus* in Northern Peninsular Malaysia and Its Potential Epidemiological Implications. PLoS ONE 2010; 5(7): e11790. doi:10.1371/journal.pone.0011790

19. Malaysia Reports 130,000 Dengue Cases In 2019, Highest Since 2015. Available from: https://codeblue.galencentre.org/2020/01/03/malaysia-reports-130000-dengue-cases-in-2019-highest-since-2015/ Accessed 3 May 2020

20. iDengue. Available from: http://idengue.arsm.gov.my/ Accessed 22 June 2020

21. Anker M, Arima Y. Male-female differences in the number of reported incident dengue fever cases in six Asian countries. Western Pac Surveill Response J, 2011; 2(2):17–23.

22. Ziegler T.; Mamahit A.; Cox N.J. 65 years of influenza surveillance by a World Health Organization-coordinated global network. Influenza Other Resp. Viruses 2018, 12, 558–565.

23. Prime Minister’s Office of Malaysia. Coronavirus Disease 2019 (Covid-19). Available from: https://www.pmo.gov.my/special-contents/2019-novel-coronavirus-2019-ncov/ Accessed 22 April 2020

24. Trash in Penang drops by almost 20%. Available from: https://www.thestar.com.my/news/nation/2020/03/26/trash-in-penang-drops-by-almost-20 Accessed 30 April 2020

25. Only eight dengue hotspots in Penang. Available from: https://www.thestar.com.my/news/community/2005/10/27/only-eight-dengue-hotspots-in-penang Accessed 2 April 2020

26. MCO: Travel limited to 10km from home. Available from: https://www.malaysiakini.com/news/518198 Accessed 30 April 2020

27. Service MW. A critical review of procedures for sampling populations of adult mosquitos. Bull Entomol Res. 1977;67:343–82.

28. Scott Wand Morrison AC. Vector Dynamics and Transmission of Dengue Virus: Implications for Dengue Surveillance and Prevention Strategies Vector Dynamics and Dengue Prevention. Current topics in microbiology and immunology 2010; 338:115–28

29. Mark H Myer, Chelsea M Fizer, Kenneth R Mcpherson, Anne C Neale, Andrew N Pilant, Arturo Rodriguez, Pai-Yei Whung, John M Johnston, Mapping *Aedes aegypti* (Diptera: Culicidae) and *Aedes albopictus* Vector Mosquito Distribution in Brownsville, TX. Journal of Medical Entomology. 2020 Jan; 57,231–240, https://doi.org/10.1093/jme/tjz132

30. Smith N. Worst dengue outbreak for seven years in Singapore linked to coronavirus lockdown Available from: https://www.telegraph.co.uk/global-health/science-and-disease/worst-dengue-outbreak-seven-years-singapore-linked-coronavirus/ Accessed 30 April 2020

31. Jindal A and Rao S. Lockdowns to Contain COVID-19 Increase Risk and Severity of Mosquito-Borne Disease Outbreaks. medRxiv: doi: https://doi.org/10.1101/2020.04.11.20061143 [Preprint]. 2020 [cited 2020 June 2]. Available from: https://www.medrxiv.org/content/10.1101/2020.04.11.20061143v1.full.pdf+html

32. Harrington LC, Thomas W Scott, Kriangkrai Lerdthusnee et al. Dispersal of the dengue vector *Aedes aegypti* within and between rural communities. The American journal of tropical medicine and hygiene. March 2005; 72(2):209–20

33. Rahim J, Ahmad AH, Maimusa AH and Irfan Shah. Updated abundance and distribution of *Aedes albopictus* (Skuse) (Diptera: Culicidae) in Penang Island, Malaysia Tropical Biomedicine 2018; 35(2): 308

34. World Health Organization (WHO). Dengue increase likely during rainy season: WHO warns. Available from: https://www.who.int/westernpacific/news/detail/11-06-2019-dengue-increase-likely-during-rainy-season-who-warns Accessed 30 April 2020

35. Scott TW and Morrison AC. *Aedes aegypti* density and the risk of dengue-virus transmission Vol2 Ecological Aspects for Application of Genetically Modified Mosquitoes ISBN: 978-1-4020-1585-4

36. Nur Aida H, Abu Hassan A, Nurita AT, Che Salmah MR, Norasmah B. Population analysis of *Aedes albopictus* (Skuse) (Diptera:Culicidae) under uncontrolled laboratory conditions. Trop Biomed 2008; 25: 117–125

37. Bonizzoni M, Gasperi G, Chen X, and James AA. The invasive mosquito species *Aedes albopictus*: current knowledge and future perspectives. Trends in parasitology, 2013; 29(9), 460–468. https://doi.org/10.1016/j.pt.2013.07.003

38. Dengue control – The Mosquito. Available from: https://www.who.int/denguecontrol/mosquito/en/ Accessed 22 June 2020

